# Perfusable Apparatus For Thick-tissue Creation And Growth (patch) Of Cardiac Tissue

**DOI:** 10.1101/2022.07.18.500065

**Authors:** Isaree Pitaktong, Yusheng Jason He, Katherine Nurminsky, Tyler Dunn, Amatullah Mir, Sarah Koljaka, Olivia Dunne, Stephanie Ran, Wesley Shih, Anya Wang, Hiroshi Matsushita, Daniel Rodgers, Narutoshi Hibino

**Affiliations:** Section of Cardiac Surgery, Department of Surgery, University of Chicago, 5841 S. Maryland Ave. Chicago, IL 60637, USA; Pediatric Cardiac Surgery, Advocate Children’s Hospital, 4440 W 95th St. Oak Lawn, IL 60453, USA

## Abstract

Cardiac tissue engineering has been developed as a potential alternative treatment for heart failure. However, current 3D tissues are limited in size and thickness due to the lack of an effective vascularization method. We have developed a novel bioreactor system to create viable vascularized cardiac tissue from multicellular spheroids using a digital light processing (DLP) 3D bioprinting system. Spheroids were created from induced pluripotent stem cells (iPSC) and cardiac fibroblasts (FB) using special dimple plates for mass production. One centimeter cubic tissues were created from spheroids using a DLP 3D printed mold with vascular channels. The tissue was maintained in a perfusion chamber under regulated flow and pressure following differentiation to cardiac tissue and endothelialization. Mass production of large spheroids (35,000 / tissue, diameter of 395.99 um +/- 101.15 um) was achieved from 170 million iPSCs and 50 million FBs for the creation of 1cm3 cardiac tissue in a 3D printed mold with vascular channels. The cardiac tissues (n=5) were perfused for 20 days under stable pressure of 17.5 +/- 3.05 PSI and flow of 5000 uL/min +/- 1116.42 uL/min. On days 10 and 20, Alamar blue assays showed viability for all five tissues (Alamar blue intensity: Day 10 1.57 +/- 0.15. Day 20 2.21 +/- 0.19). Thick and viable cardiac tissues were created and maintained using a 3D printed vascularized mold and perfusion system for maturation and growth in vitro for 30 days. This technology will open new doors for viable in vitro cardiac tissue creation.

## Introduction

Heart failure contributes to approximately 287,000 deaths a year (Emory). The recent rise of 3D bioprinting in the medical field has offered several clinical applications to heart failure treatment. Printing artificial valves and other anatomically relevant structures that can allow for more efficient treatment of heart failure and cardiovascular disease. Printed cardiovascular tissues contain great advantages in preoperative decisions and intraoperative navigation. 3D printing is gradually changing the common traditions of treatment and diagnosis for patients with heart failure, cardiovascular disease, and other heart-related illnesses (Wang, 2020). In the human body, related cells that are joined together are collectively referred to as tissue, and these cells work together as organs to accomplish specific functions in the human body. Blood vessels around the cells vascularize, providing nutrients to the tissue to keep it healthy. The vascularized, thick-tissue models will function as organ analogs, or models. These will be used to study the mechanism of disease or develop new drugs. To properly construct a 3D printed tissue, factors such as vascularization, perfusion and circulation, and cell viability must be considered in order to ensure the maintenance and proper function of the cardiac patch (Puluca, 2019).

Current literature on cardiac tissue perfusion contains missing information on perfusion and vascularization (Kato, 2021). Since it is difficult to create thick tissue because of vascularization, more careful methodologies are being developed to ensure that cell-to-cell contact is maintained to promote vascularization. While existing methodologies tend to mix spheroids and cells in the ink prior to printing (Maiullari, 2018), our approach uses spheroids to promote vascularization. In our approach, we investigate perfusion and vascularization through the development of a 1 cm^3^ 3D printed cardiac tissue mold using hydrogel bioink. Beating, vascularized cardiac tissue from multicellular spheroids consisting of human-induced pluripotent stem cells (hiPSC) derived cardiomyocyte, fibroblasts (FB), and endothelial cells (EC) were grown using the mold in order to ensure proper vascularization in the cardiac patch. Our method used spheroids to promote vascularization. Multiple analyses were conducted throughout the experiment to observe tissue volume, cell viability, and differentiation. Following these analyses, an in vitro perfusion system was used to perfuse the cardiac tissue for 30 days, and tissue viability was measured in order to determine the functionality of the printed tissue. Analysis throughout the 30 day period was performed by tracking pressure and flow of the perfusion system.

## Methods

### Mixed spheroid creation using large dimple plate

Human-induced pluripotent stem cells (hiPSC) and cardiac fibroblasts (FB) were collected from separate monolayer cultures and joined in large dimple plates for the creation of large-scale mixed spheroids (**Figure 1**). To make the large dimple plates, 100% EtOH is added to a large dimple plate and left to soak overnight, then washed away. The plate is autoclaved, placed in a petri dish, and a solution of cell media and cell solution is combined and added to the plate. The plate is placed on a shaker for multiple days. After spheroids form, the plate is decannulated and the spheroids are transferred to a 50 mL conical tube. The hiPSCs and FBs were combined in a 70:30 ratio respectively. The mixed cell suspension was dispensed into a large dimple plate (Tissue by Net Ltd., Tokyo, Japan) for mixed spheroids formation in Essential 8 Medium (A1517001 Invitrogen) for four days before being collected and transferred to a 3D printed mold. A total of 10 mL was contained in each dimple plate, composed of 3.9 × 10^6^ hiPSCs, 1.7 × 10^6^ FBs, and the remaining volume composed of E8 medium (which was dependent on cell solution concentration). A total of 32 large dimple plates, including 125 million hiPSCs and 55 million FBs were used for the creation of a 1cm^3^ patch.

**Figure 1:**
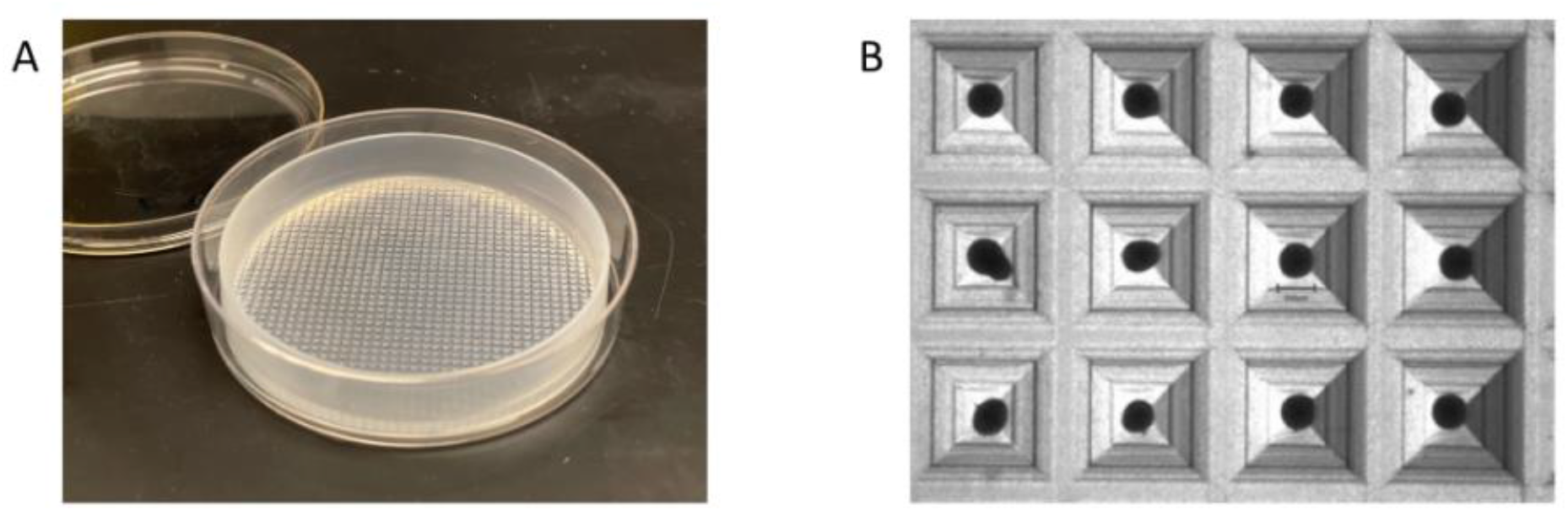
Spheroid creation in large dimple plate. Picture of large dimple plate which cells are seeded into and form spheroids (A). Spheroid formation in each hole of dimple plate at 1.25x [2] (B).

### Hydrogel creation using 3D printer

The 3D printer consists of a 405 nm industrial projector reflected upward toward the photoink. The photoink, Polyethylene glycol diacrylate (PEGDA) 575 g/mol, and gelatin methacrylate (GelMA) were mixed in a 1 to 1 ratio, total volume 2 mL, and placed on the PDMS-lined Vat directly between the UV light projection and building platform. At this point, a stereolithographic (STL) file, detailing the geometry of our mold, was uploaded to the printer. Specifications including light intensity and exposure time were set based on the specifications of the photoink manufacturer, 20 mW/cm^2^ and 5 seconds. After the printer reads the file and the parameters for the print set, the build platform lowers into the photoink droplet and builds in 100 micrometers layers. This 3D printed mold has 16 vascular channels, arranged in a 4 × 4 configuration, which allowed culture medium to flow through the tissue unidirectionally. The internal space volume of the mold was 1 cm^3^ (**Figure 2A**).

**Figure 2:**
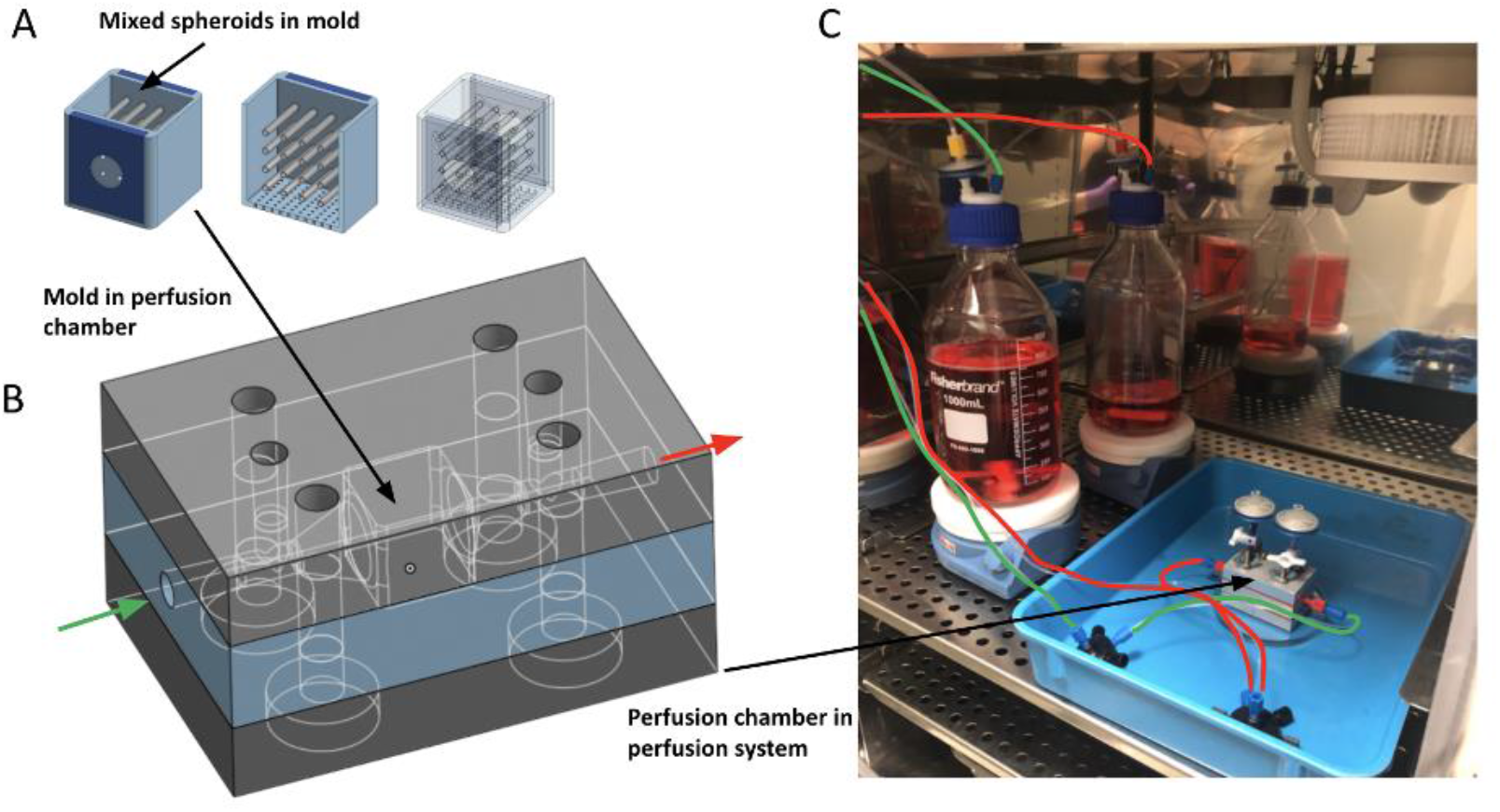
Perfusion chamber. The 3D printed mold (1cm^3^) with sagittal cuts depicting vascular channels and top-down view of channels to show patency filled with mixed spheroids (A). 3D printed mold placed inside of perfusion chamber where media flows unidirectionally (B). The green and red arrows describe the flow within the perfusion system. The mold was placed in the center of the chamber, as described by the black arrows. The perfusion chamber was connected to our perfusion system which included two 1000 mL media reservoirs, a perfusion controller, and compressed air tanks to push media from an inlet media reservoir through the mold, and into an outlet media reservoir (C). A rotary valve between the media reservoirs and perfusion chamber alternates inlet and outlet reservoirs upon completion of 1000 mL perfusion to keep the flow through perfusion chamber unidirectional (not pictured).

### Creation and perfusion of 1cm^3^ cardiac tissue

After the addition of spheroids to our 3D printed mold (**Figure 2A**), the spheroid-filled mold was placed into the perfusion chamber, a 3-tier metal device that allowed for media to flow through the patch (**Figure 2B**). The tissue must include channels composed of endothelial cells with active medium perfusion throughout the tissue for the duration of the trial. Medium perfusion must be enough to keep the tissue surviving and there must be histological evidence that demonstrates fluid flow in the vascular channels. Once seeded into the mold and connected with the perfusion system (**Figure 2C**), the differentiation of hiPSCs to cardiomyocytes (CM) was started by temporal modulation of Wnt signaling using small molecules (CHIR99021, Tocris, R&D Systems, Cat. No. 4423, and IWR-1, Sigma-Aldrich, Cat. No. I0161). The E8 medium was switched out for B27(-)/CHIR. After another 48 hours, the medium was switched again, this time with just B27(-). The following day, the B27(-) medium was switched for B27(-)/IWR-1. And after two additional days, the medium was switched for the last time with B27(+)/EGM. At this point, the vascular channels of the 3D printed mold were endothelialized by flowing human umbilical vein endothelial cells (HUVEC). The completion of endothelialization was set as day 0. B27(+)/EGM medium was perfused for the following 30 days.

The perfusion system was propelled by compressed air flowing through a regulator. The regulator was set to keep flow under strict 3.5 PSI and 1 mL/min flow rate constraints. For each additional patch added to the perfusion system in parallel, PSI and flow rate were increased by 3.5 and 1 mL/min respectively. The 3D printed mold allowed us to use vascular channels, 500 microns in diameter, to flow medium through the tissue.

### Evaluation methods

#### Viability Assay

Cell viability of the 3D printed cardiac tissue was determined using Alamar Blue assay. Alamar blue is a non-toxic and cell-permeable reagent. Resazurin is the active ingredient in the reagent and upon entering living cells, the cell environment reduces resazurin to resorufin, which is red and highly fluorescent. Because only viable cells with active metabolism can reduce resazurin to resorufin, a change in fluorescence and color of the media surrounding cells exposed to Alamar blue is indicative of living cells. The change in fluorescence (and therefore cell viability) can be detected with the naked eye, but quantified via plate reader absorbance.

#### Histology

Tissues were fixed in 4% paraformaldehyde, embedded in histogel then paraffin, and cut into 5um sections. The tissues were then washed in PBS, incubated in 0.5% Triton X for 5-10 minutes, washed in PBS again, and blocked for one hour using 10% goat serum. Primary antibodies to the cell protein or biomarker of interest (Troponin T (cardiomyocyte), Vimentin (fibroblast), CD31 (endothelial cells), Oct-4 (iPSC)) are incubated with the sections overnight. Removal of unbound primary antibodies was accomplished by washing using PBS. Secondary antibodies were incubated with the sections for one hour before washing using PBS and mounting using Vectashield Antifade Mounting Medium with 4’,6-diamidino-2-phenylindole (DAPI) (Vector Laboratories, Burlingame, CA).

#### Perfusion Record

Perfusion data including flow and pressure were automatically and continuously recorded in the perfusion pump system.

## Results

### Tissue volume analysis

The volume of vascular channels inside of the 3D printed mold was 142.732mm^3^. The space for cells to filled in the 1cm^3^ cubic mold (internal volume = 1000mm^3^) mold was (1000-142.732)/1000 = 0.857. The mixed cell spheroids were able to fill more than 85% of 1cm cubic tissue when the tissue was placed in the perfusion system. Total 5 tissues were created for different follow-up dates, day 10 (n=1), day 20 (n=1), and day 30 (n=3, named as day 30-1, 30-2, 30-3). At each time point, the mold was collected from the perfusion chamber and the size of the tissue inside of the mold was measured as shown in (**Figure 3**). The ratio of the tissue size at each time point compared with the original tissue prior to placing in the perfusion chamber was 52.5% at day 10, 90.7% at day 20, 73.8%±14.5% at day 30 (30-1: 60.9%, 30-2: 71.1%, 30-3: 89.6%). The day 10 tissue showed significantly low volume due to mechanical damage by high perfusion pressure when only this patch was placed in the system to begin the trial. Following the initiation of day 10 tissue perfusion, 4 other patches were added and a total of 5 patches were perfused together. Therefore, the pressure for the other 4 patches was not as high as the day 10 tissue, which enabled proper maintenance of the tissue. On day 30, as some molds began to break down, sample tissues were damaged when removed from the perfusion chambers, which led to lower volume compared with day 20. The precision of the 3D printer made it possible to evenly perfuse the patch throughout the 30-day period (Figure 2A).

**Figure 3:**
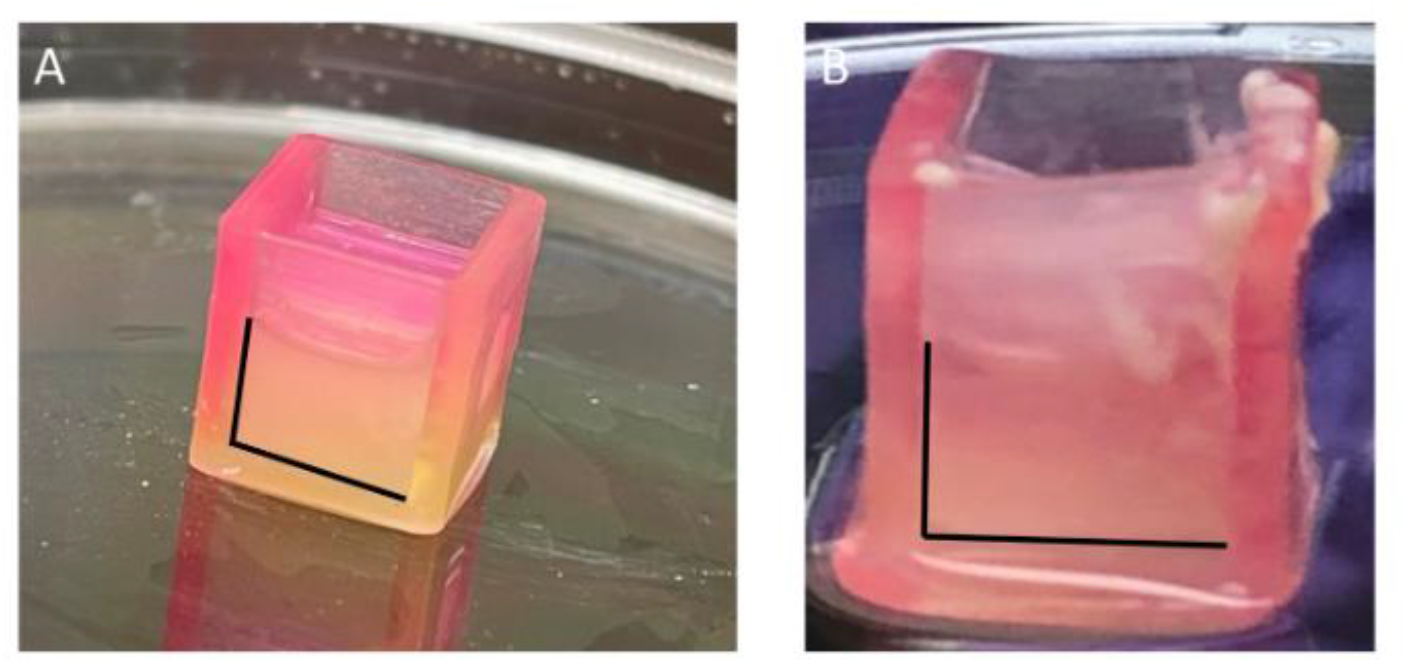
Tissues created in the 3D printed mold. The size of the tissues were 52.5% patch 1 (not pictured), 90.7% patch 2 (A), 73.8%±14.5% patch 3 (not pictured), 71.1% patch 4 (not pictured), and 89.6% at patch 5 (B). Day 10 tissue, patch 1, was damaged due to initial high-pressure perfusion. The day 30 tissues, patches 3-5, showed lower volume than day 20, patch 2 because the sample tissues were damaged due to broken mold when they were taken out from the perfusion chambers.

### Viability assay

Viability assays were conducted on days 10, 20, and 30 for each patch, keeping the tissues in their perfusion chambers. The assays were conducted using Alamar blue, perfused through one patch at a time in a simple circuit running at approximately 1 mL/min. The Day 10 average absorbance (ratio of sample/HUVEC control) was 1.57 ± 0.15. On Day 20, the average absorbance was 2.21 ± 0.19. Day 30 had an average absorbance of 2.63 ± 0.05 (**Figure 4**). Overall, there was a marked increase compared with the HUVEC control, which had an absorbance of 1.273. This control was made to ensure that cells within the spheroids were viable and metabolizing the assay alongside the HUVECs that lined the channels.

**Figure 4:**
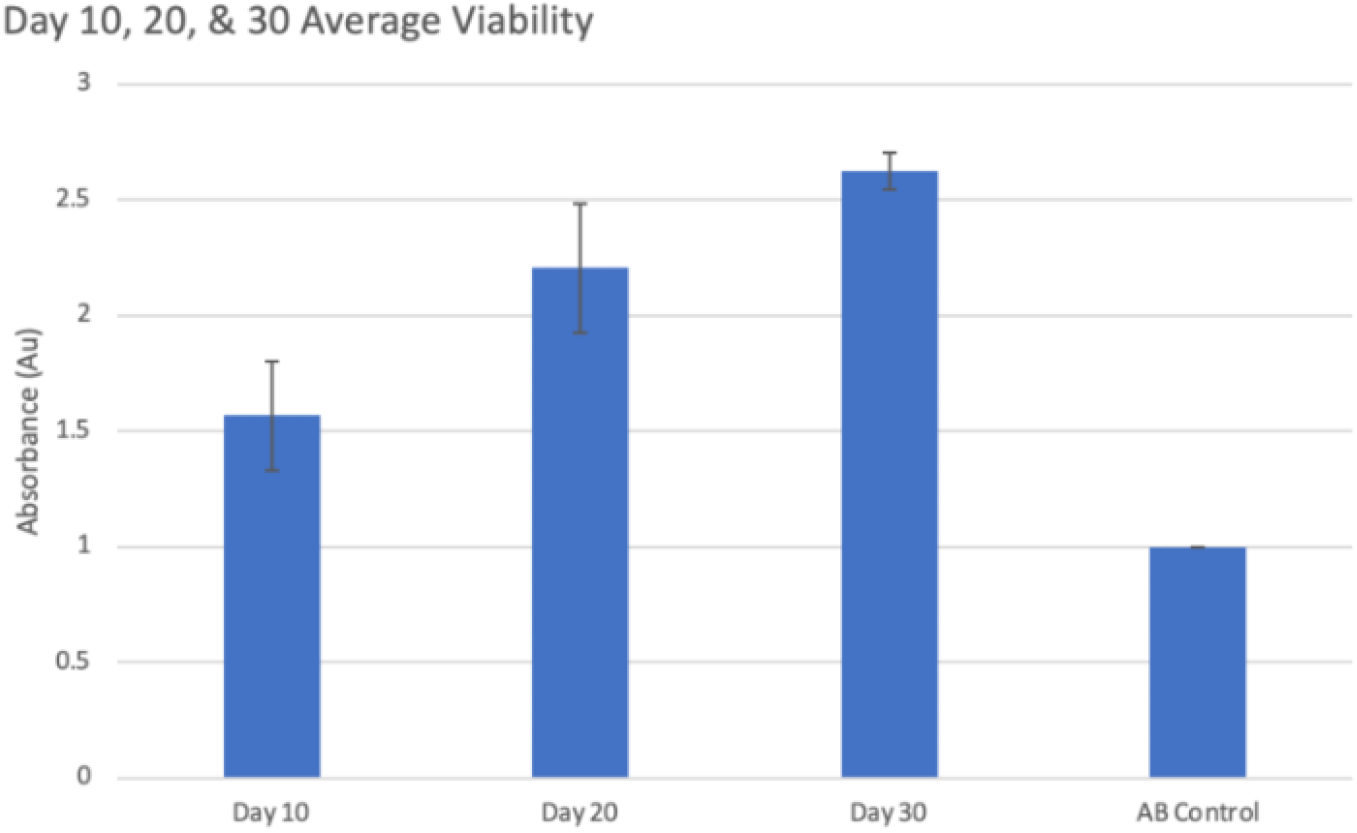
Alamar blue assay. Average values for Day 10, 20, and 30 % transmittance ratio with HUVEC control.

### Histology

Immunofluorescence of the tissues showed differentiated cardiomyocytes from iPSC (Oct 4 negative (A) and Troponin-T positive (C)), endothelial cells lined up on the lumen of vascular channels (B), and fibroblasts (D) (**Figure 5**).

**Figure 5:**
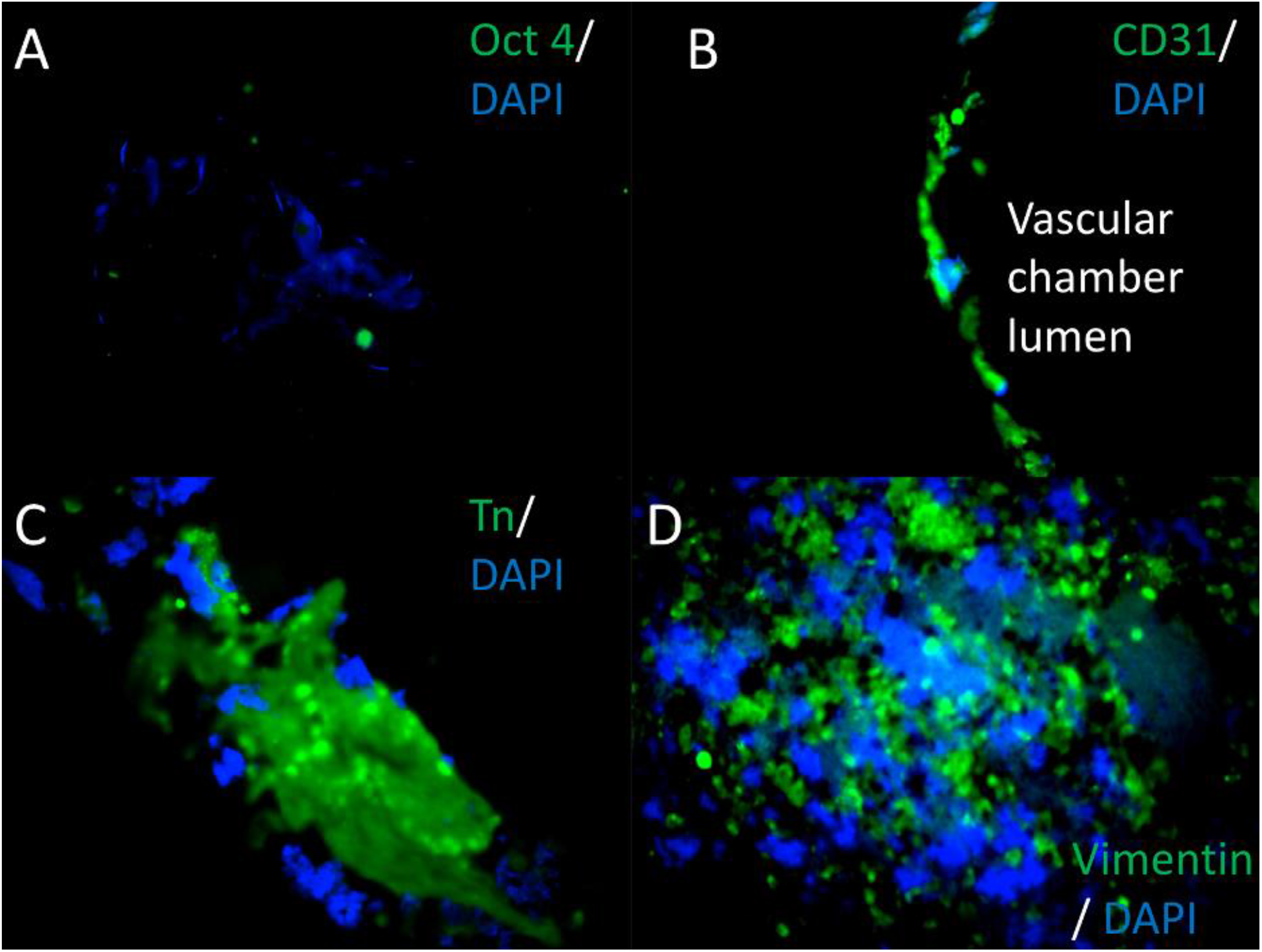
Histology. Histological analysis of tissues.

**Figure 6:**
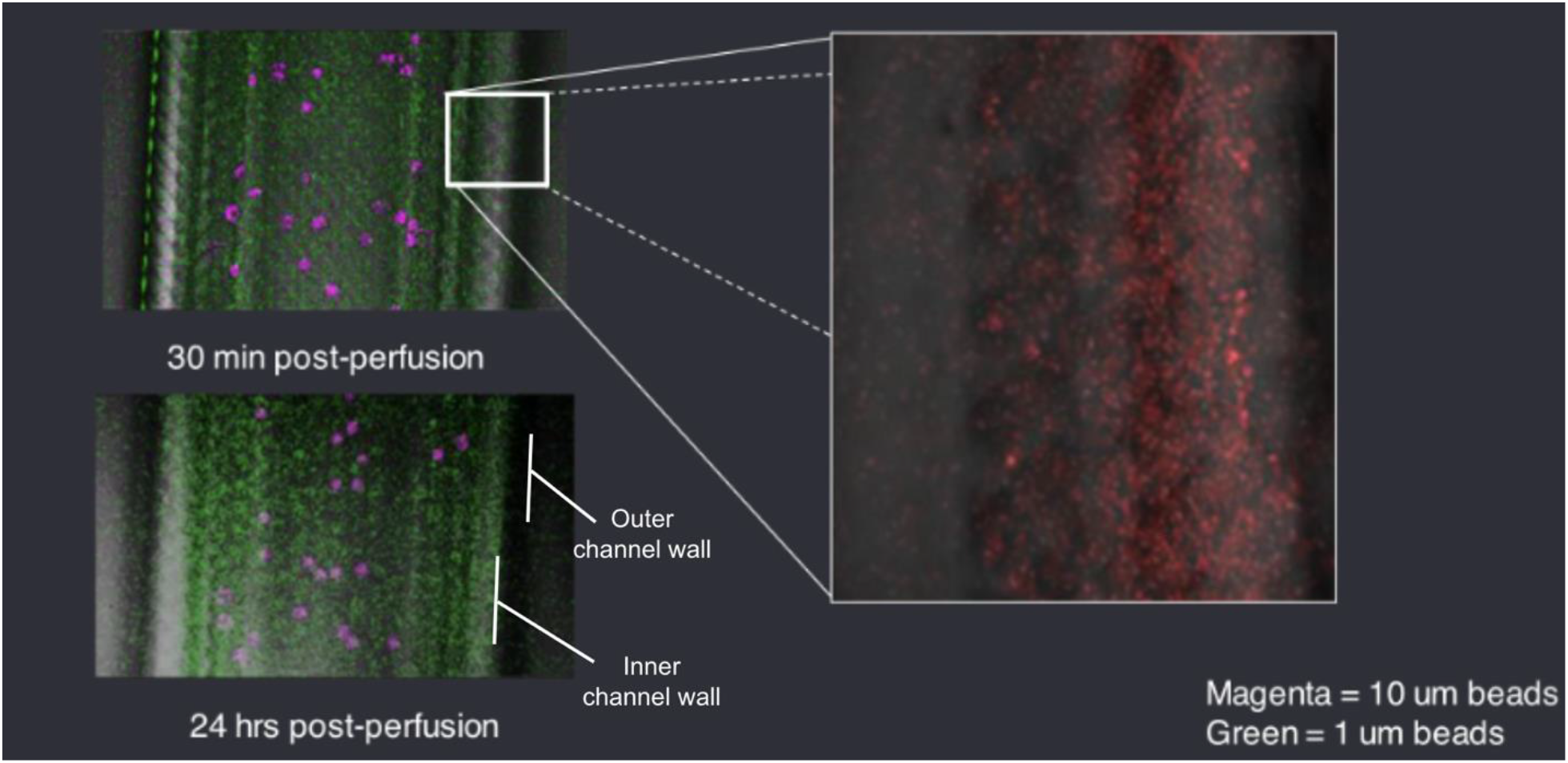
Perfusion. Post perfusion images.

**Figure 7:**
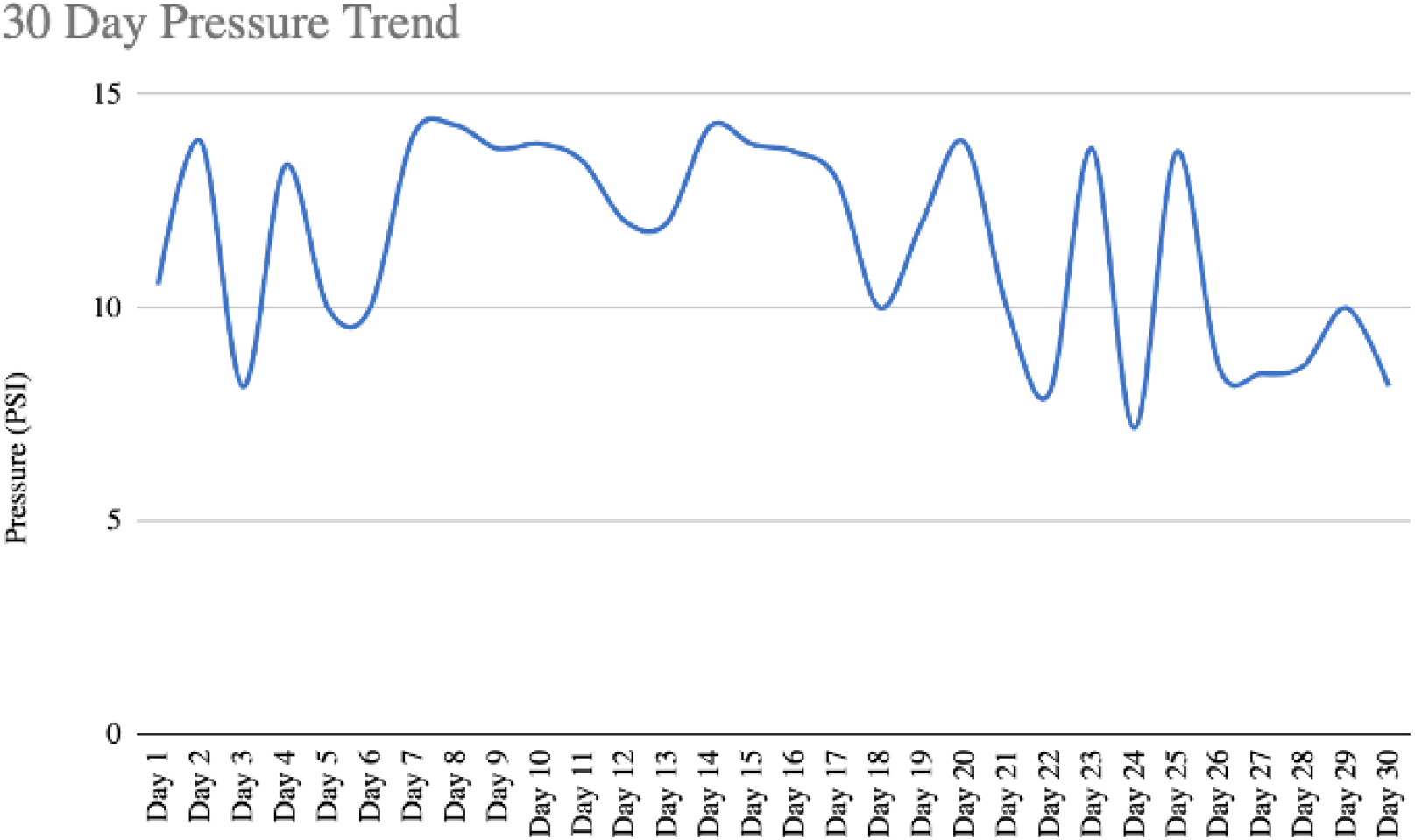
Perfusion system pressure. Due to the inability of accurate liquid flow-rate control, PSI was maintained for 30-days.

**Figure 8:**
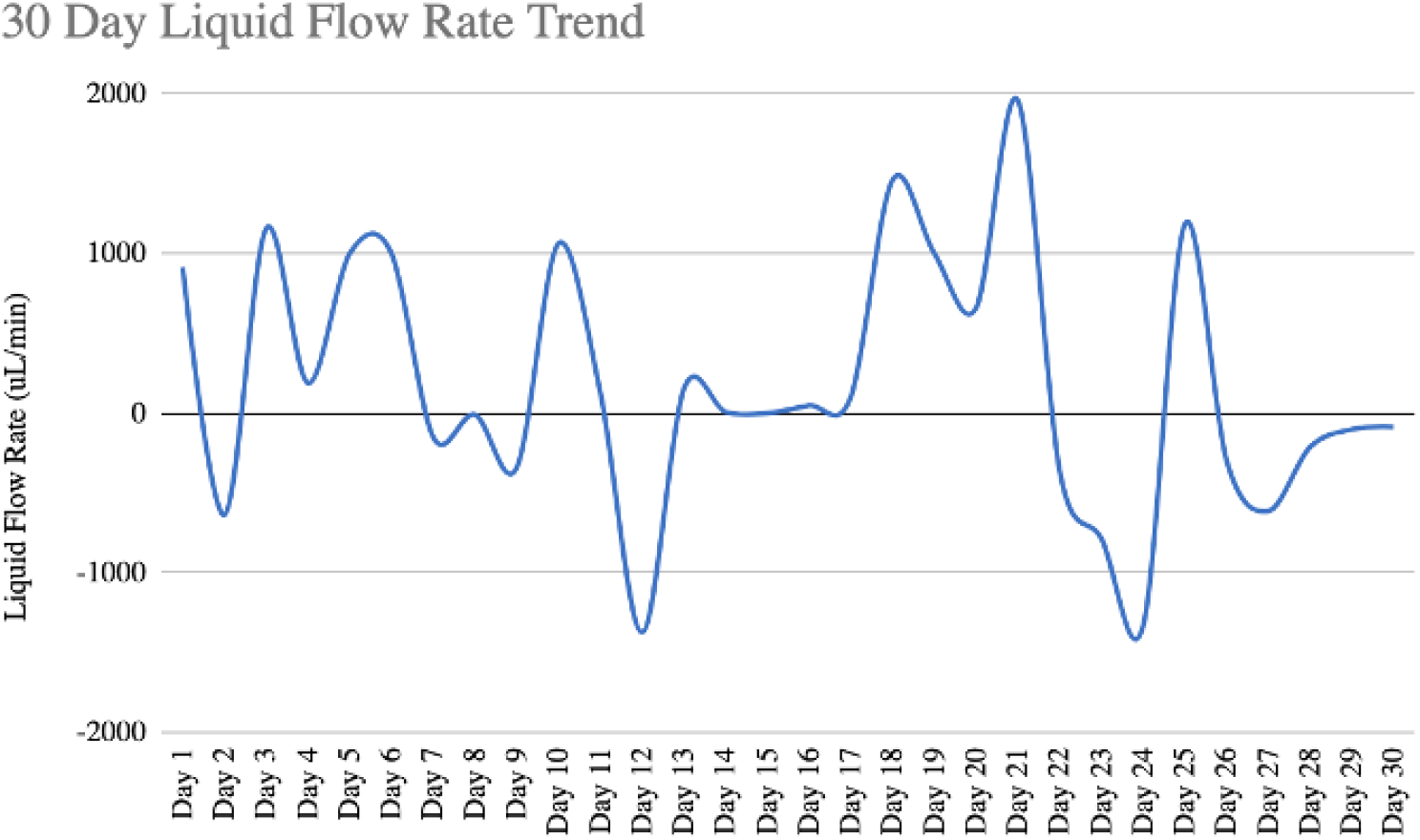
Perfusion system liquid flow rate. Negative flow rate values depict changes in flow direction due to pressure alterations in the system.

### Perfusion analysis

To evaluate the function of vascular channels, we perfused 10um and 1um magnetic micro particles based on polystyrene through the vascular channels for 30 min. The large diameter of the particles in the dye did not allow for diffusion from vascular channels suggesting there was no leakage in the perfusion system during the 30-day trial.

The continuous record of flow and pressure in the perfusion system was tracked and recorded for the entirety of the 30-day trial. The change of pressure and flow throughout the first 6 days of patch creation was due to the manipulation of culture medium for differentiation and human umbilical vein endothelial cells (HUVEC) perfusion. The negative readings for flow rate after the first 6 days were due to changes in flow direction as the medium was changed. Constant pressure suggested continuous perfusion without interruption for 30 days.

## Discussion

The results obtained from this study demonstrate important advances in the field of in vitro heart tissue generation. Firstly, the vascular channel perfusion yielded continuous perfusion according to the constant pressure and flow observed. The viability analysis showed a steady increase in average absorbance compared to the control HUVEC sample. This result is a strong indication of maintaining cell viability within the spheroids lining the vascular channel. The findings of this study indicate that the 30 day perfusion trial with the mold yielded a maintaining cell viability.

The balance between parenchymal tissue and vasculature is necessary in the development of functional 3D tissues. Recent studies focus on the creation of vasculature. For instance, Kinstlinger et. al utilizes selective laser sintered printing and GelMA in the development of their molds (Kinstlinger, 2020). They developed complex dendritic networks that maintained shape after perfusion. Kinstlinger et al’s study uses these networks in their experiment, in contrast to the 16 linear channels used in our experiment. While this method improves the ability to build vasculature and branching, their method does not leave much space for parenchymal and cardiomyocyte cells. Maiullari et. al’s vascularization study developed a sophisticated microfluidic-based printing head (MPH) for the simultaneous extrusion of multiple bioinks (Maiullari, 2018). In contrast to this experiment which did not mix cells with bioink, they instead printed a patch with two separate bioinks with the MPH, which yielded better differentiation and more functional cardiomyocyte organization as a result of the spatial orientation within the printed fibers. To promote vascularization support, Maiullari et. al’s experiment used intercalated endothelial precursor cells within the 3D cardiac structure to generate the Janus patch (which alternated HUVEC and iPSC cells between layers) and two other patches, the 4:2:4 and 2:2:2:2:2 layered patches. Ameliorated vascularization was observed in the multi-cellular bioprinted structures in comparison to the controls that did not contain endothelial cells. While Maiullari et. al’s method is prints with endothelial cells mixed with the bioink, our method uses spheroids to enhance vascularization in the patch. Changing the printing method to print multi-cellular bioprinted patches and using spheroids (as done so in our method) may allow for vascularization. Both Kinstlinger’s and Maiullari’s methods rely on bioprinting patterns for vasculature design. However, natural vasculature is much more complicated. Our method is a combination of both the bioprinting pattern and natural vascularization from spheroids. Our method has the advantage of maintaining an organized vasculature creation that allows for the balance between parenchymal tissue and vasculature within the mold.

The flow pattern and speed are important variables for tissue maturation in flow chamber system. We used high constant flow for perfusion, but Homan’s study used various flow. Homan et. al perform an in vitro flow vascularization and maturation of kidney organoids. They develop a millifluidic culture system in order to determine the effects of the extracellular matrix, media composition, fluidic shear stress (FSS), and culture with human endothelial cells on in vitro development of kidney organoids (Homan, 2019). Our experiment, which did not vary FSS in its perfusion of the cardiac patch, vastly differs from in the methods of perfusion compared to their experiment. They subject the organoids to superfusion overnight over a range of various fluid rates, with varying FSS. After 10 days, they observed formation of vascular networks in organoids. In our experiment, instead of strictly increasing flow rate, we can perfuse the patch at higher rates of FSS in order to enhance vascularization in the mold. Cardiac tissue is in pulsatile flow, and the effect of various flow on tissue formation should be investigated in future studies.

As a limitation of this study, while the histology, perfusion system, and tissue volume analyses of the cardiac patch showed strong cell viability, decreased tissue volume, and proper perfusion capabilities, our study does not include the analysis of tissue efficacy. This study also did not demonstrate contractability and conductivity properties of the mold. For future research, the findings of this study can be extended by examining larger tissue sizes greater than 1 cm3 in volume. Testing various bioinks such as natural polymers or synthetic polymers should also be explored in order to determine the most durable and conductive mold. The conductivity and contractility of the mold should also be investigated in future studies.

## Conclusion

We created 1 cm^3^ vascularized cardiac tissue from multicellular spheroids consisting of human induced pluripotent stem cell (hiPSC) derived cardiomyocytes, fibroblasts, and endothelial cells using a 3D printed hydrogel mold. We did not mix the cardiomyocytes with biomaterial to maintain cell-cell contact which was the critical component for functional cardiac tissue. We demonstrated the feasibility to perfuse the cardiac tissue while maintaining tissue viability in our unique perfusion system for 30 days. The volume of the tissue was slightly decreased after 30 days. there were Troponin-T positive cardiac cells in the tissue and CD31 positive vascular channels, suggesting some function of cardiac tissue. The mechanical property, detailed structure of vascular network, actual function such as contractability and electrical conductivity will be evaluated in the future.

